# Endothelial sphingosine 1-phosphate receptors promote vascular normalization to influence tumor growth and metastasis

**DOI:** 10.1101/606434

**Authors:** Andreane Cartier, Tani Leigh, Catherine H. Liu, Timothy Hla

## Abstract

Sphingosine 1-phosphate receptor-1 (S1PR1) is essential for embryonic vascular development and maturation. In the adult, it is a key regulator of vascular barrier function and inflammatory processes. Its roles in tumor angiogenesis, tumor growth and metastasis are not well understood. In this report, we show that S1PR1 is expressed and active in tumor vessels. Tumor vessels that lack S1PR1 (*S1pr1* ECKO) show excessive vascular sprouting and branching, decreased barrier function, and poor perfusion accompanied by loose attachment of pericytes. Compound knockout of *S1pr1*, 2 and 3 genes further exacerbated these phenotypes, suggesting compensatory function of endothelial S1PR2 and 3 in the absence of S1PR1. On the other hand, tumor vessels with high expression of S1PR1 (*S1pr1* ECTG) show less branching, tortuosity and enhanced pericyte coverage. Larger tumors and enhanced lung metastasis were seen in *S1pr1* ECKO whereas *S1pr1* ECTG showed smaller tumors and reduced metastasis. Furthermore, anti-tumor activity of doxorubicin was more effective in *S1pr1* ECTG than the wild-type counterparts. These data suggest that tumor endothelial S1PR1 induces vascular normalization and influences tumor growth, evolution and spread. Strategies to enhance S1PR1 signaling in tumor vessels may be an important adjunct to standard cancer therapy.

**Significance:** Endothelial sphingosine 1-phosphate receptors modulate tumor angiogenesis by inducing vascular normalization, which allows better blood circulation and enhanced anti-tumor therapeutic efficacy.

## Introduction

Sphingosine-1-phosphate (S1P), a lysophospholipid found in blood and lymph, regulates cell survival, migration, immune cell trafficking, angiogenesis, and vascular barrier function. S1P binds to a family of G protein-coupled sphingosine 1-phosphate receptors 1 to 5 (S1PR1-5) which are expressed on most cells (1). The prototypical S1PR1, which is abundantly expressed in vascular endothelial cells (EC), is required for embryonic vascular development and maturation (2, 3). S1PR1 inhibits VEGF-induced vascular sprouting (4) by promoting interactions between VE-Cadherin and VEGFR2 that suppress VEGF signaling (5). However, S1PR1 function is compensated by other S1PRs that are expressed in EC, albeit at lower levels. For example, S1PR2 and S1PR3, which are both capable of signaling via the Gi pathway, function redundantly as S1PR1 in embryonic vascular development (6). Mice that lack S1PR1, 2 and 3 exhibit early embryonic lethality similar to global (7) or red blood cell-specific (8) sphingosine kinase (SPHK)-1 and −2 double knockout mice that lack circulatory S1P. These findings support the notion that coordinated signaling of VEGF-A via its receptor tyrosine kinases and plasma S1P via EC G protein-coupled S1PRs is fundamental for the development of a normal primary vascular network.

Tumor progression requires new vessel growth, a phenomenon termed as tumor angiogenesis. This is achieved by the production of angiogenic factors which activate endothelial cells from pre-existing blood vessels to undergo angiogenesis (9). For example, angiogenic stimulators such as VEGF-A are released by tumor cells to induce angiogenesis and tumor growth (10). Angiogenesis is also associated with spreading of tumors to metastatic sites. Tumor vessels, characterized by abnormal morphology, are highly dysfunctional in their barrier and transport properties (11). Strategies to induce phenotypic change of tumor vessels to resemble normal vessels, termed vascular normalization, has been attempted (11-13). Indeed, anti-VEGF antibodies induce vascular normalization in preclinical models and in the clinic, which may in part explain their efficacy in the treatment of metastatic cancer. After anti-VEGF treatment, tumor vessels show increased perfusion and efficacy of anti-tumor chemotherapies. However, preclinical studies have shown a precise time window for the efficacy of antiangiogenic therapies, as prolonged antiangiogenic treatment can lead to excessive pruning, hypoxia, activation of alternative proangiogenic pathways and the development of resistance (14).

Even though S1P signaling via endothelial S1PRs is a central player in vascular development, the role of S1P signaling axis in tumor angiogenesis and progression is not clear. Early studies showed that S1PR1 is expressed in tumor vessels and downregulation of its expression with 3’UTR-targeted siRNAs suppressed tumor growth (15). However, administration of FTY720, a prodrug that is phosphorylated and binds to 4 out of 5 S1P receptors, suppressed tumor growth and metastasis in mouse models (16, 17). Application of VEGF pathway inhibitors together with S1PR-targeted small molecules achieved better inhibition of tumor angiogenesis (18). However, precise roles of endothelial S1PR subtypes in tumor angiogenesis, progression and metastasis has not been analyzed in genetic models. We systematically studied mouse genetic models in which S1PRs have been modified either alone or in combination, and studied tumor vascular phenotypes in syngeneic lung cancer and melanoma models. We show that endothelial S1PRs are key regulators of vascular normalization and that stimulation of this pathway enhances chemotherapeutic efficacy.

## Results

### S1PR1 expression and signaling regulates tumor vascular phenotype

S1PR1 is expressed in angiogenic vessels of tumors grown subcutaneously in mice (15). In order to determine whether S1PR1 is actively signaling in angiogenic endothelial cells, we used a mouse model referred to as S1PR1-GFP signaling mice that allows visualization of the β-arrestin recruitment to S1PR1 (19). We injected Lewis lung carcinoma cells (LLC) subcutaneously in S1PR1-GFP signaling mice and analyzed the resected tumor sections by fluorescence microscopy. GFP positivity was observed in tumor vessel like structures. GFP^+^ cells were co-localized with PECAM-1, but not α-smooth muscle actin (Fig. 1A-D). These data suggest that S1PR1 signaling is active in endothelial cells of angiogenic tumors.

**Figure 1.**
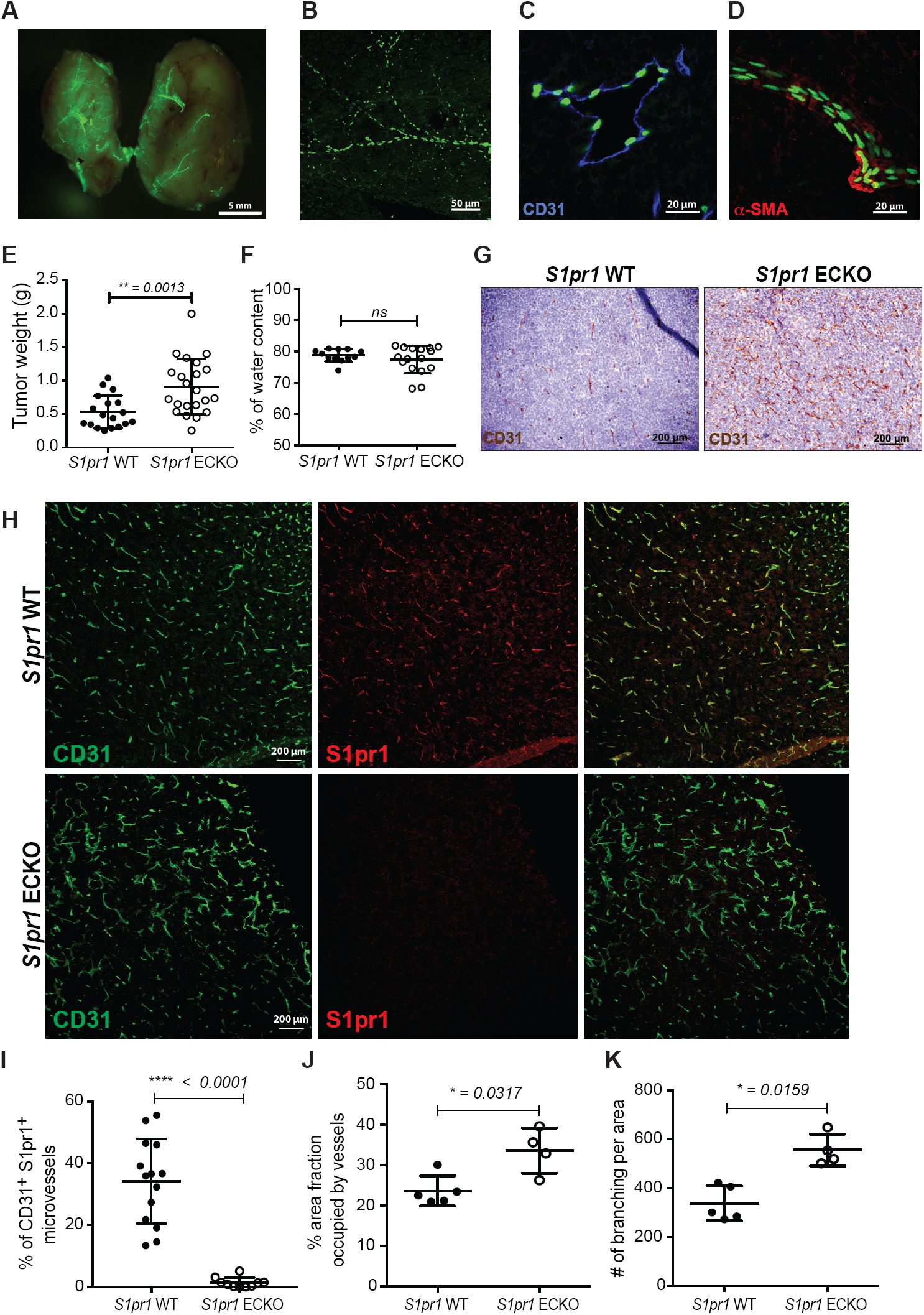
Loss of EC-specific S1PR1 induces tumor growth and angiogenesis. (A-D) Subcutaneous LLC tumors grown in S1P1 GFP signaling mice show positive GFP signal in vascular structures, as confirmed by (A) whole mount fluorescence imaging, (B-D) two-photon and confocal microscopy of 35 µm tumor sections stained with PECAM-1 (CD31) (C) and *α*-SMA (D). *n*=3 independent experiments. (E) Weight of subcutaneous LLC tumors grown in control *S1pr1* WT and ECKO mice, and quantification of their water content (F). *n*=6 independent experiments containing 3 or 4 mice per group, expressed as a mean of 2 tumors per animal ± SD. (G) Immunohistochemistry (IHC) image of 12 µm paraffin sections stained with CD31 antibody show tumor vascular density in *S1pr1* WT and ECKO mice by light microscopy. Representative images from 2 separate experiments containing 4 mice per group are shown. (H-I) Immunofluorescence (IF) of 35 µm sections of tumor, frozen in OCT and stained with S1PR1 and CD31 antibodies, shows extensive deletion of S1PR1 signal in endothelial cells by confocal microscopy. Double positive S1PR1^+^CD31^+^ signal was also quantified. Images are representative of 2 separate experiments containing 4 mice each. (J-K) Quantification of vascular density and branching from immunofluorescence images (*n*=10-14 (I) and 4-5 (J-K)) from tumors grown in control *S1pr1* WT and ECKO mice. Data are expressed as mean ± SD. *P* values were determined by two-tailed unpaired Mann-Whitney test comparing control *S1pr1* WT and *S1pr1* ECKO mice. *ns*, nonsignificant, *P*\* ≤ 0.0332, *P*\*\* ≤ 0.0021, *P*\*\*\* ≤ 0.0002, *P*\*\*\*\*≤ 0.0001.

To assess the functional role of S1PR1 in tumor angiogenesis, we used a mouse model in which *S1pr1* is deleted specifically in endothelial cells by tamoxifen-activated Cre recombinase (*S1pr1*^*flox/flox*^ *Cdh5 Cre-ER*^*T2*^), which is referred to as *S1pr1* ECKO (4, 20-22). Tumors grown in *S1pr1* ECKO mice were almost 2 times bigger than the tumors grown in control mice (Fig. 1E). Water content in both the *S1pr1* ECKO and control mice was similar (Fig. 1F), suggesting that increased vascular leak in the tumor is not the cause of the increase in tumor size. Histological analysis did not reveal marked changes in matrix accumulation. These data suggest that increased tumor cell proliferation and/or recruitment of host-derived cells may be the reason for increased tumor size.

To determine the functional role of S1PR1 in tumor angiogenesis, vascular density and morphology were assessed in tumor sections and analyzed by light and confocal fluorescence microscopy followed by quantitative image analysis (Fig. 1G-I). Tumor vessels in *S1pr1* ECKO mice show higher vascular density, excessive sprouting and branching (Fig. 1G-K). These data indicate that S1PR1 is present and active in tumor vascular endothelial cells and suppresses hypersprouting of intratumoral vessels.

### Loss of S1PR1 expression in endothelial cells impairs mural cell coverage of tumor vessels

During embryonic development, S1PR1 expression on endothelial cells is important for vascular stabilization and mural cell recruitment (3). Sections from tumors grown in *S1pr1* ECKO and control animals were assessed for mural cell coverage. Immunohistochemical staining with alpha smooth muscle actin (*α*-SMA) antibody show that tumor vessels from *S1pr1* ECKO are deficient in smooth muscle actin positive cell coverage (Fig. 2A-C). On the other hand, NG2^+^ pericytes were similar in number in both WT and *S1pr1* ECKO tumor vessels (Fig. 2D,E). However, pericyte attachment to the endothelial cells in the tumor vessels from *S1pr1* ECKO mice appeared loose, with pericytes weakly adhered to the endothelial cell layer, which is in sharp contrast to the WT counterparts. The lack of *α*-SMA+ mural cell coverage, and loose association of NG2^+^ pericytes, may in part explain the altered vascular morphology seen *in S1pr1* ECKO tumor vessels.

**Figure 2.**
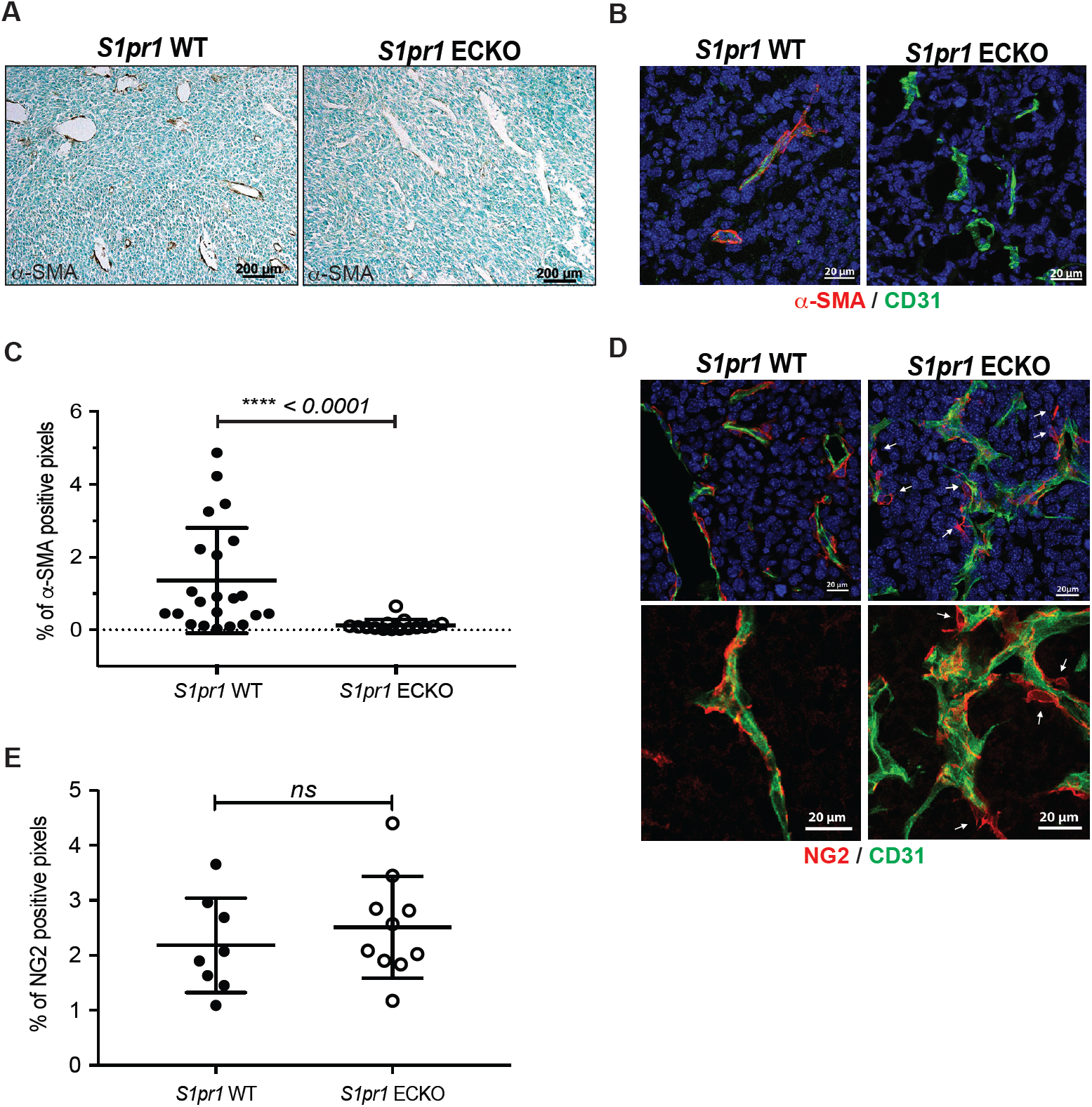
Loss of EC-specific S1PR1 impairs mural cell coverage. (A-C) IHC (A) and IF (B) of tumor sections from *S1pr1* WT and ECKO mice and stained with *α*-SMA antibody and CD31. (C) Quantification of *α*-SMA positive pixels from confocal images (*n*=15-22) of sections from tumors grown in control *S1pr1* WT and ECKO mice. (D) Low and high magnification confocal images of frozen OCT tumor sections stained with CD31 and pericyte marker NG2. Arrows indicates area of loose pericyte coverage of the endothelium. (E) Quantification of total NG2 positive signal from confocal images (*n*=8-10) of sections from tumors grown in control *S1pr1* WT and ECKO mice. Data are expressed as mean ± SD. *P* values were determined by two-tailed unpaired Mann-Whitney test comparing control *S1pr1* WT and *S1pr1* ECKO mice. ns, nonsignificant, *P*\*\*\*\*≤ 0.0001.

### Tumors in *S1pr1* ECKO mice show increased vascular permeability and metastatic potential

Since tumor vessels from *S1pr1* ECKO mice showed deficient maturation, we characterized their vascular barrier properties. We performed intravenous injection of high molecular weight fluorescein isothiocyanate conjugated (FITC)-dextran (2000 kDa) and tetramethylrhodamine-dextran (70 kDa) and assessed vascular leak in both tumor vessels and in the lung. Quantification of tissue sections from tumors and lung tissue from *S1pr1* ECKO show increased leakage of the 70 kDa dextran in the proximity of the vessels (Fig. 3A-D) while the 2000 kDa dextran clearly delineated tumor vascular lumens.

**Figure 3.**
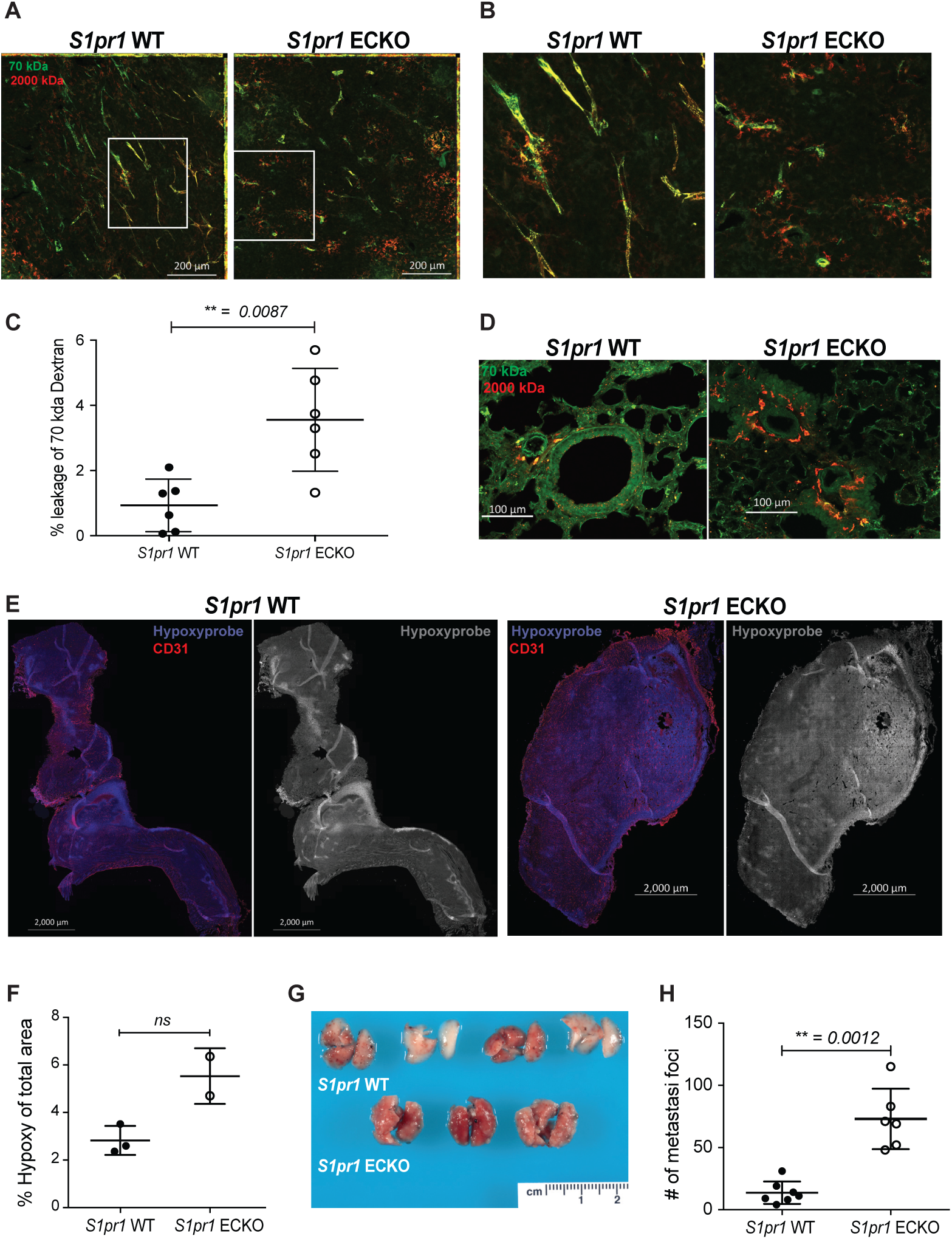
EC-specific S1PR1 deficient tumor vessels show increased leakage. Tiled (A) and zoomed (B) confocal images of 35 µm tumors sections grown in *S1pr1* WT and ECKO mice, intravenously injected with 70 kDa (Fluorescein, green) and 2000 kDa (TMR, red) dextran. (*n*=6 per group) (C) Quantification of extravasated 70kDa dextran from confocal images (*n*=6). Data are expressed as mean ± SD. *P* values were determined by two-tailed unpaired Mann-Whitney test comparing control *S1pr1* WT and *S1pr1* ECKO mice. *P* ≤ 0.05 (D) Confocal images of lung sections from tumor bearing *S1pr1* WT and ECKO mice that were injected intravenously with 70 kDa (Fluorescein, green) and 2000 kDa (TMR, red) dextran (*n*=6 per group). (E-F) Tumor hypoxia in *S1pr1* WT and ECKO mice was visualized by Hypoxyprobe-1 injected intravenously and IF staining with Hypoxyprobe-1 antibody. Confocal images of whole tumors and quantification of Hypoxyprobe-1 staining images (*n*=3). *P* values were determined by two-tailed unpaired *t* test with Welch’s correction comparing control *S1pr1* WT and *S1pr1* ECKO mice. *ns*, nonsignificant (G) Lung colonization of B16-F10 cells injected intravenously in *S1pr1* WT and ECKO mice. Image is representative of 3 independent experiments. (H) Quantification of metastasis foci. Data are expressed as mean ± SD. *P* values were determined by two-tailed unpaired Mann-Whitney test comparing control and *S1pr1* ECKO mice. *P*\*\* ≤ 0.0021

Since vascular leakage could lead to tissue perfusion defects and hypoxia (23), we used the oxygen-sensitive probe termed as hypoxyprobe-1 to determine the hypoxia status of tumors in WT and *S1pr1* ECKO mice (24). As shown in Fig. 3E and F, hypoxic zones of *S1pr1* ECKO tumors trended towards an increase even though the difference was statistically not significant, suggesting a minor change in tumor oxygenation.

Intravenous tail vein injection of B16F10 melanoma cells into WT and *S1pr1* ECKO mice, which results in lung metastasis, showed markedly increased metastatic nodules in the lungs of the mice that lack endothelial S1PR1 (Fig. 3G,H), suggesting that aforementioned vascular defects contributed to lung colonization of circulating tumor cells and metastasis.

Taken together, these data show that S1PR1 expressed on endothelial cells regulates tumor angiogenesis, vessel maturation, vascular permeability, and tumor perfusion thus influencing primary tumor growth and metastatic potential.

### Redundant functions of S1PR2 and 3 in the regulation of the tumor vascular phenotypes, tumor growth and metastasis

Endothelial cells express S1PR2 and S1PR3 in addition to S1PR1 (25). While S1PR1 and S1PR2 induce opposing cellular effects, for example, in barrier function, S1PR2 can activate redundant signaling pathways in the absence of S1PR1 (26-28). In addition, both S1PR2 and S1PR3 are capable of signaling redundantly as S1PR1; for example, via the G_i_ pathway (29-32). The roles of these receptors in tumor angiogenesis has not been examined.

We recently developed a conditional mutant allele for *S1pr2* and developed a mouse model for *S1pr2* ECKO using the tamoxifen-inducible Cdh5-Cre driver (33). Using this mouse model with LLC tumors, angiogenesis tumor vascular phenotypes were analyzed. As shown in Fig. 4A, tumors grown in *S1pr2* ECKO mice were significantly smaller than those in the WT counterparts. Tumor vasculature showed no significant changes in permeability to intravenously injected 70kD fluorescent dextran (Fig. 4B). However, increased pericyte coverage was seen (Fig. 4C). Moreover, intravenously injected B16 melanoma cells showed decreased metastatic potential in the lungs of *S1pr2* ECKO (Fig. 4D). These results reveal the opposing functions of S1PR1 and S1PR2 in tumor vascular phenotype regulation.

**Figure 4.**
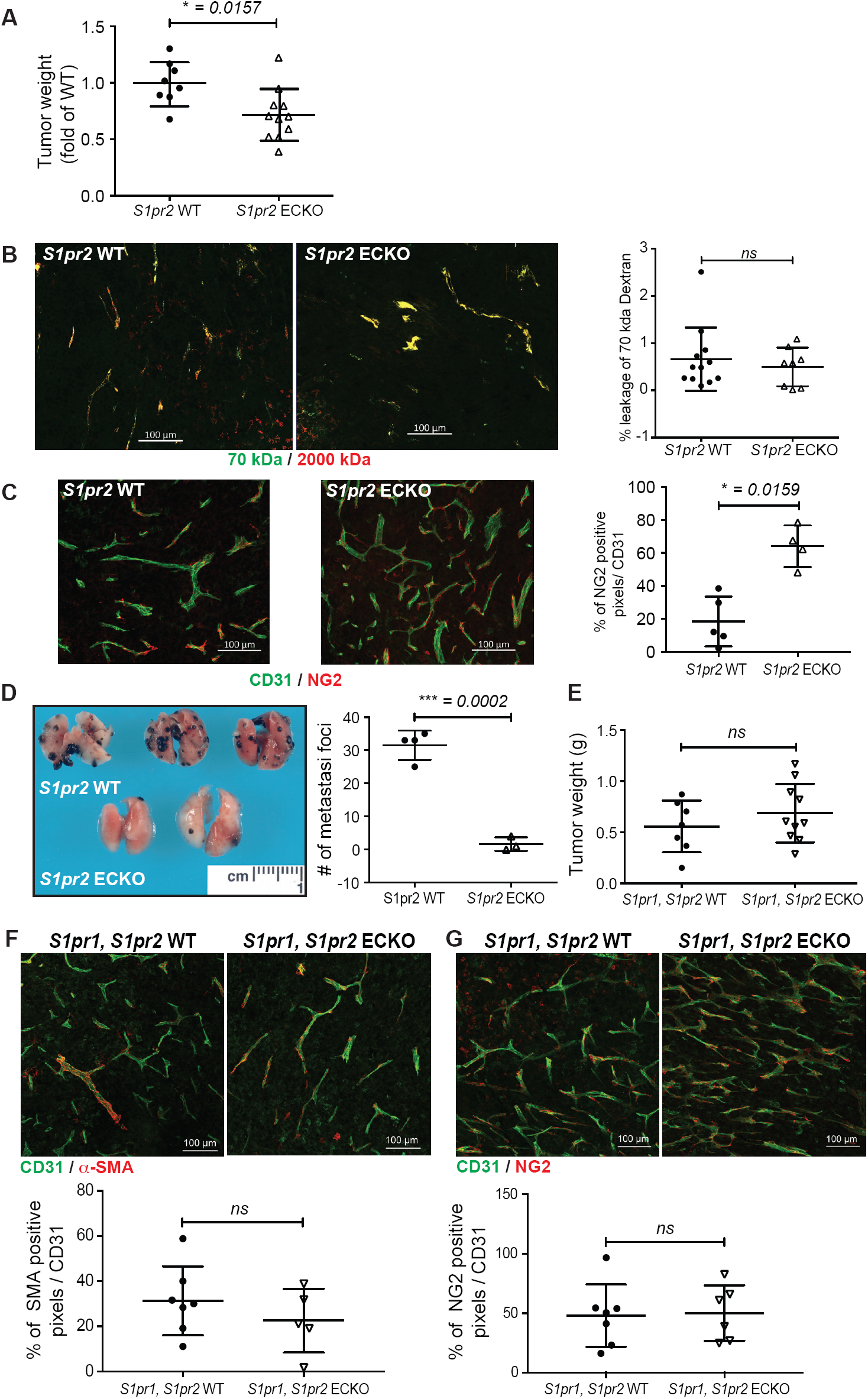
Loss of EC-specific S1PR2 impairs tumor growth and metastasis. (A) Whole tumor weight of subcutaneous LLC tumors grown in *S1pr2* WT and ECKO mice. *n*=3 independent experiments containing 3 or 4 mice per group, expressed as a mean of 2 tumors per animal ± SD. (B) Tiled confocal images of 35 µm tumors sections grown in *S1pr2* WT and ECKO mice, intravenously injected with 70 kDa (Fluorescein, green) and 2000 kDa (TMR, red) dextran. (*n*=3 per group) (C) Quantification of extravasated 70kDa dextran from confocal images (*n*=8-12 per group). Data are expressed as mean ± SD. *P* values were determined by two-tailed unpaired Mann-Whitney test comparing control *S1pr2* WT and *S1pr2* ECKO mice. (C) Confocal images of tumor sections stained with CD31 and pericyte marker NG2 and quantification of total NG2 positive signal from confocal images (*n*=8-10) of sections from tumors grown in control *S1pr2* WT and ECKO mice (n=4-5 per group). *P* values were determined by two-tailed unpaired Mann-Whitney test comparing control *S1pr2* WT and *S1pr2* ECKO mice. *P*\* ≤ 0.0332. (D) Lung colonization of B16-F10 cells injected intravenously in *S1pr2* WT and ECKO mice. Image is representative of 3 independent experiments. Quantification of metastasis foci is expressed as mean ± SD. *P* values were determined by two-tailed unpaired *t* test with Welch’comparing control and *S1pr2* ECKO mice. *P*\*\*\* ≤ 0.0002. (E) Whole tumor weight of subcutaneous LLC tumors grown in *S1pr1, S1pr2* WT and ECKO mice. *n*=3 independent experiments containing 3 mice per group, expressed as a mean of 2 tumors per animal ± SD. *P* values were determined by two-tailed unpaired Mann-Whitney test comparing *S1pr1, S1pr2* WT and ECKO mice. *ns*, nonsignificant. (F) 35µm tumor sections from *S1pr1, S1pr2* WT and ECKO mice were stained with CD31 and *α*-SMA or pericyte marker NG2 and show vascular morphology and mural cell coverage. Quantification from confocal images (n=5-7 per group) of total *α*-SMA and NG2 positive signal per CD31 positive signal are expressed as mean ± SD. *P* values were determined by two-tailed unpaired Mann-Whitney test comparing control and *S1pr2* ECKO mice. *ns*, nonsignificant.

When compound *S1pr1 S1pr2* ECKO mice were analyzed, subcutaneous LLC tumors were similar in size, and NG2^+^ and *α*-SMA^+^ mural cell recruitment to tumor vessels was not different from the control counterparts (Fig. 4E). Tumor vascular phenotype showed modest hypersprouting (Fig. 4F-G), suggesting that the effects of *S1pr1* ECKO were neutralized by the lack of S1PR2, which mediates opposite endothelial phenotypic effects. Additionally, the percent of *α*-SMA^+^, CD31^+^ and NG2^+^, CD31^+^ double signal were similar in both *S1pr1 S1pr2* WT and ECKO mice (Fig.4F-G).

We next examined the redundant role of S1PR3 in tumor angiogenesis. When *S1pr3*^*-/-*^ mice (6) were compared with compound *S1pr1* ECKO *S1pr3*^*-/-*^ mice, tumor growth, vascular density and the recruitment of *α*-SMA^+^ and NG2^+^ mural cells to tumor vessels largely resembled that of *S1pr1* ECKO mice (Fig. 5A-E). However, compound triple KO of *S1pr1* and *S1pr2 ECKO* in *S1pr3*^*-/-*^ background showed marked increase in tumor growth, vascular hypersprouting and mural cell disengagement phenotypes (Fig. 5F,G). These data suggest that S1PR3 functions are redundant to S1PR1 in suppressing endothelial hypersprouting as well as properties of highly abnormal vascular phenotypes. Together, these findings support the redundant functions of S1PR2 and S1PR3, which compensate the function of attenuated S1PR1.

**Figure 5.**
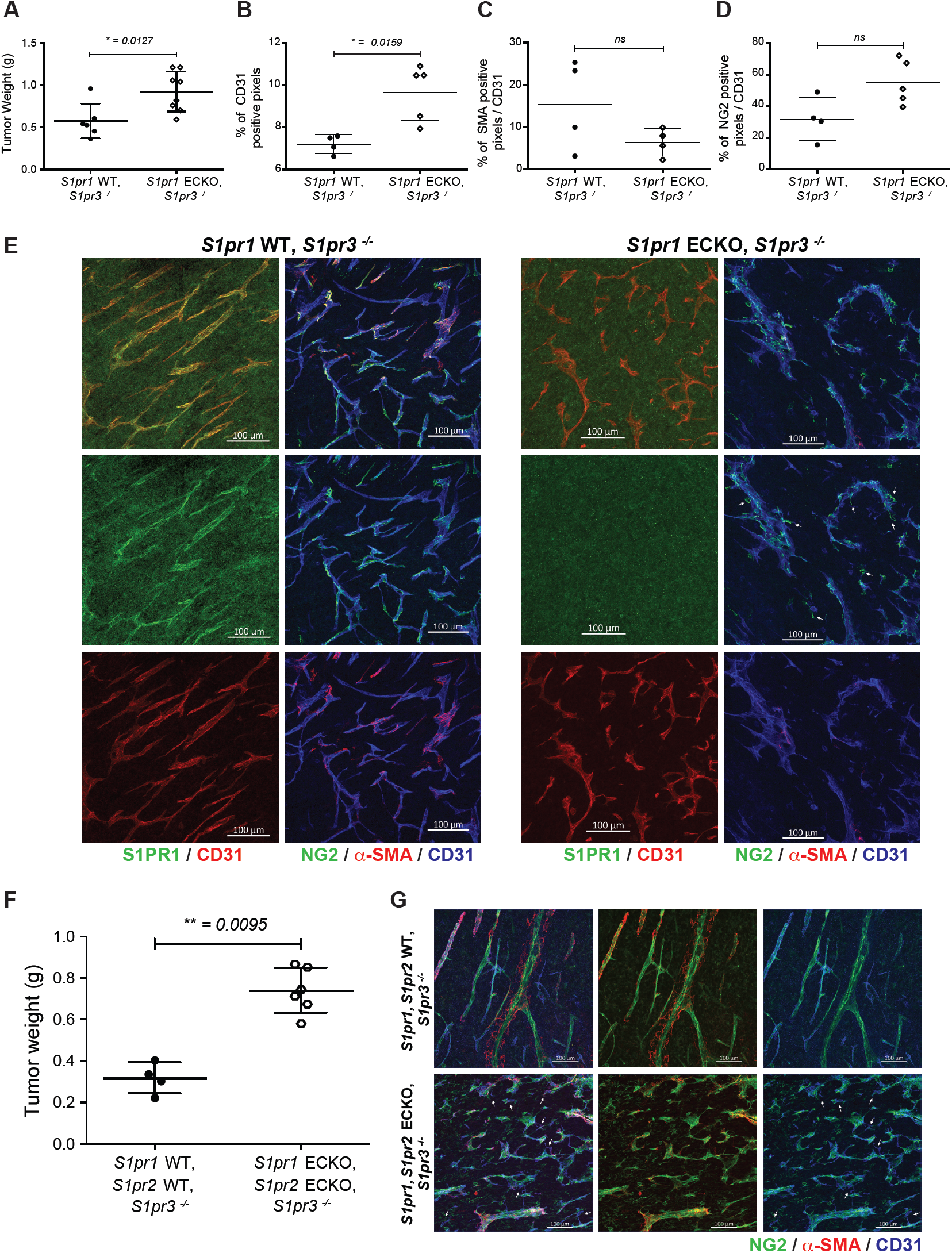
Compound endothelial-specific deletion of *S1pr1* and *S1pr2* in *S1pr3*^*-/-*^background induces tumor growth and severe vascular disorganization. (A) Whole tumor weight of subcutaneous LLC tumors grown in *S1pr1* WT, *S1pr3*^-/-^ and *S1pr1* ECKO, *S1pr3*^*-*/-^ mice. *n*=2 independent experiments containing 3 or 4 mice per group, expressed as a mean of 2 tumors per animal ± SD. Quantification of images (n=4-5) of 35 µm tumor section show the percentage of CD31 signal (B), the *α*-SMA^+^ – CD31^+^(C) or CD31^+^ - NG2^+^ (D) positive signal. *P* values were determined by two-tailed unpaired Mann-Whitney test comparing *S1pr1* WT, *S1pr3*^-/-^ and *S1pr1* ECKO, *S1pr3*^*-*/-^ mice. *ns*, nonsignificant, *P*\* ≤ 0.0332 (E) Immunofluorescence (IF) of 35 µm sections of tumor grown in *S1pr1* WT, *S1pr3*^-/-^ and *S1pr1* ECKO, *S1pr3*^*-*/-^ mice and stained with S1PR1, CD31, NG2 and *α*-SMA antibodies, shows extensive deletion of S1PR1 signal in endothelial cells by confocal microscopy, vascular morphology and mural cell coverage. (F) Whole tumor weight of subcutaneous LLC tumors grown in *S1pr1* WT, *S1pr2* WT, *S1pr3*^-/-^ and *S1pr1* ECKO, *S1pr2* ECKO, *S1pr3*^*-*/-^ mice. *n*=2 independent experiments containing 2 or 3 mice per group, expressed as a mean of 2 tumors per animal ± SD. *P* values were determined by two-tailed unpaired Mann-Whitney test comparing *S1pr3*^-/-^ and *S1pr1* ECKO, *S1pr2* ECKO, *S1pr3*^*-*/-^ mice. *P* ≤ 0.05 (G) 35µm tumor sections stained with CD31, *α*-SMA and NG2 antibodies show vascular morphology and mural cell coverage.

### Overexpression of S1PR1 in endothelial cells promotes normalization of tumor vessels and enhances chemotherapeutic efficacy

Due to the prominent role of endothelial S1PR1 in tumor vasculature, growth and metastatic potential, we used the inducible S1PR1 endothelial-specific transgenic mice (*S1pr1*^*flox/stop/flox*^ *Cdh5 Cre-ER*^*T2*^) (ECTG) (4, 20). Subcutaneous LLC tumor size was smaller in *S1pr1* ECTG mice (Fig. 6A) while water content was similar (Fig. 6B). Overexpression of S1PR1 in tumor vessels (Fig. 6C-E) showed less vascular branches and sprouts characterized by more linear and less tortuous vascular morphology (Fig. 6F-G), which is in marked contrast to the *S1pr1* ECKO counterparts described above.

**Figure 6.**
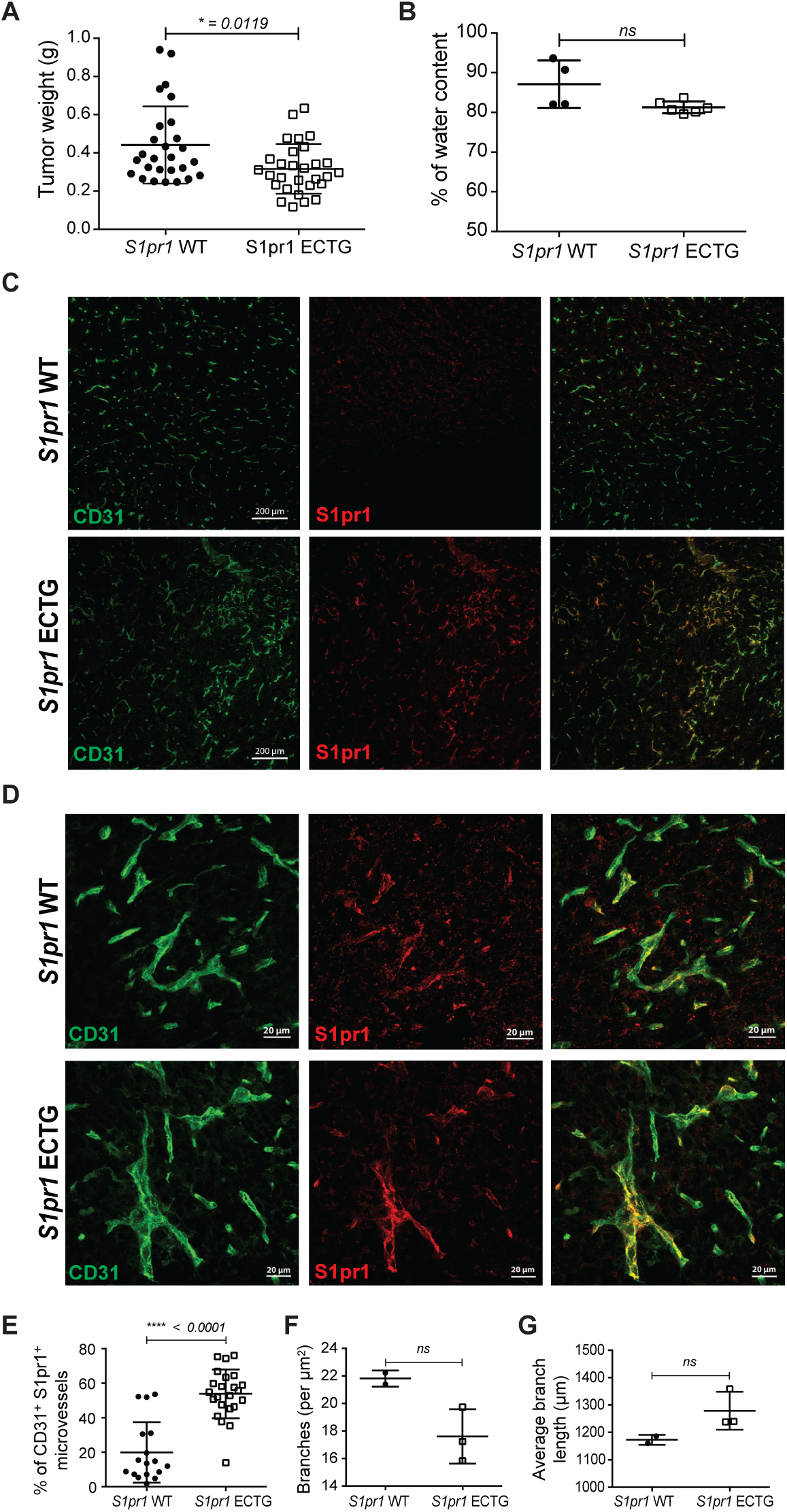
Over expression of EC-specific S1PR1 reduces tumor growth and angiogenesis. (A) Whole tumor weight of subcutaneous LLC tumors grown in control *S1pr1* WT and ECTG mice, and quantification of their water content (B). *n*=7 independent experiments containing 3 or 4 mice per group, expressed as a mean of 2 tumors per animal ± SD. *P* values were determined by two-tailed unpaired Mann-Whitney test comparing *S1pr1* WT and ECTG mice. *P*\* ≤ 0.0332 Low (C) and high (D) magnification confocal microscopy images of 35 µm sections of tumor, and stained with S1PR1 and CD31 antibodies, shows induced expression of S1PR1 in endothelial cells and vessel morphology. Double positive S1PR1^+^CD31^+^ signal quantification (E). Quantification of vascular density and branching from immunofluorescence images (*n*=17 to 23, (F)) and average vessel length (G) from tumors grown in control *S1pr1* WT and ECTG mice. Data are expressed as mean ± SD. *P* values were determined by two-tailed unpaired Mann-Whitney test comparing control *S1pr1* WT and *S1pr1* ECTG mice. *ns*, nonsignificant, *P*\*\*\*\*≤ 0.0001.

*S1pr1* ECTG tumor vessels contained higher NG2^+^ mural cells and an increase in SMA^+^ mural cells (Fig. 7A-D). Tumor vascular leakage of intravenously-injected 70 kD dextran trended to be less than the controls but did not reach statistical significance (Fig. 7E,F). In contrast, hypoxic lesions in the tumors were markedly reduced (Fig. 7G). Intravenously injected B16F10 melanoma cells also formed fewer number of metastatic nodules in the lung (Fig. 7H,I). Together, these data suggest that increased S1PR1 expression in endothelial cells promotes tumor vascular normalization and suppresses metastatic potential.

**Figure 7.**
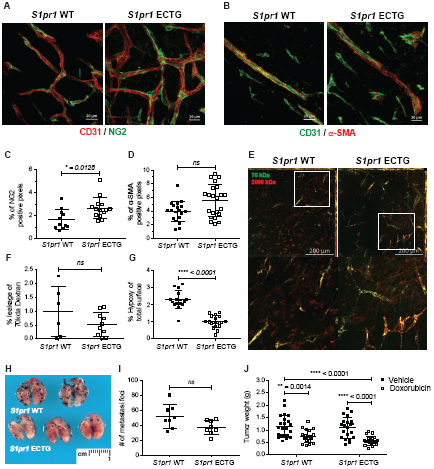
Overexpression of S1PR1 in endothelial cells enhance mural cell coverage and reduces vessel leakage. (A-B) 35 µm sections of tumor sections from *S1pr1* WT and ECTG mice and stained with NG2 or *α*-SMA and CD31 antibodies. (C) Quantification of NG2 and *α*-SMA (D) positive pixels from confocal images (*n*=11-22) of sections from tumors grown in control *S1pr1* WT and ECTG mice. Data are expressed as mean ± SD. *P* values were determined by two-tailed unpaired Mann-Whitney test comparing control *S1pr1* WT and *S1pr1* ECTG mice. *ns*, nonsignificant, *P*\* ≤ 0.0332. (E) Tiled confocal images of 35 µm tumors sections grown in *S1pr1* WT and ECTG mice, intravenously injected with 70 kDa (Fluorescein, green) and 2000 kDa (TMR, red) dextran. *n*=2 independent experiments containing 3 or 4 mice per group. (F) Quantification of extravasated 70kDa dextran from confocal images (*n*=6-10 per group). Data are expressed as mean ± SD. *P* values were determined by two-tailed unpaired ?Mann-Whitney test comparing control *S1pr1* WT and *S1pr1* ECTG mice. *ns*, nonsignificant (G) Tumor hypoxia in *S1pr1* WT and ECTG mice was visualized by Hypoxyprobe-1 injected intravenously and IF staining with Hypoxyprobe-1 antibody. Confocal images of whole tumors and quantification of Hypoxyprobe-1 staining images (*n*=2 independent experiment, and 3-4 images were quantified per group per experiment). *P* values were determined by two-tailed unpaired Mann-Whitney test comparing control and *S1pr1* ECTG mice. *ns*, nonsignificant, *P*\*\*\*\*≤ 0.0001. (H-I) Lung colonization of B16-F10 cells injected intravenously in *S1pr1* WT and ECTG mice. Image is representative of 3 independent experiments. (I) Quantification of metastasis foci. Data are expressed as mean ± SD. *P* values were determined by two-tailed unpaired Mann-Whitney test comparing control and *S1pr1* ECTG mice. (J) Whole tumor weight of subcutaneous LLC tumors grown in *S1pr1* WT and ECTG mice, treated with 5 mg/kg of doxorubicin or vehicle. *n*=8 independent experiments containing 3 or 4 mice per group, expressed as a mean of 2 tumors per animal ± SD. *P* values were determined by two-way ANOVA followed by Sidak’s multiple comparisons test comparing *S1pr1* WT and ECTG mice, ± Doxorubicin. *P*\*\*\*\*≤ 0.0001, *P*\*\* ≤ 0.0021

Since normalization of tumor vessels were shown to enhance anti-tumor therapies (11-14, 34), we injected anti-tumor chemotherapy drug doxorubicin to tumor bearing WT and S1PR1 ECTG mice every other day, starting at day 8 post injection of the LLC. As expected, doxorubicin treatment reduced the growth of the tumors in both cohorts of mice. However, in the S1PR1 ECTG mice, doxorubicin treatment was more effective in reducing tumor growth (Fig. 7J). These data suggest that S1pr1-induced tumor vascular normalization enhances the chemotherapeutic efficiency of doxorubicin.

## Discussion

S1P signaling axis via the endothelial S1PRs represent a major regulatory system for vascular maturation during development (6). Balanced signaling between angiogenic growth factors such as VEGF that signals via receptor tyrosine kinases and S1PRs which are GPCRs is essential for normal vascular development (35). In the adult, endothelial S1PR signaling regulates vascular barrier function, tone and inflammatory processes. Since tumor angiogenesis occurs postnatally, we studied the role of endothelial cell S1PR signaling axis in mouse models of tumor angiogenesis, progression and metastasis.

A principle finding of our work is that the level of S1PR expression in the tumor endothelium determines key aspects of the tumor vascular phenotype. These include endothelial sprouting, branching phenotypes and the barrier function. Lack of endothelial S1PR1 promoted excessive vascular leak, as well as markedly increased vascular sprouting and branching. We predict that attenuated S1PR1 function in the tumor endothelium would lead to decreased access of blood borne cells and substances to the tumor parenchyma. Opposite phenotypes were seen by overexpression of endothelial S1PR1. Our results suggest that S1PR1-regulated events in the newly-formed tumor vessels are important in determining their normalization status.

We also show that attenuated endothelial S1PR1 function led to increased tumor growth whereas S1PR1 overexpression led to smaller tumors. Intratumoral fluid accumulation is unable to account for the changes in tumor size. Either tumor cell proliferation and/or stromal hematopoietic cell infiltration could be affected by the signaling of endothelial S1PRs. We speculate that elaboration of angiocrine functions of tumor endothelial cells that influence tumor cells *per se* and/or cells of the tumor microenvironment are involved.

In addition to tumor vascular normalization and progression, we show that the signaling of endothelial S1PR1 influences the ability of circulating tumor cells to establish metastatic colonies in lungs. Since defective endothelial junctions leading to decreased barrier properties of the tumor vessels are controlled by this receptor, we suggest that this function of S1P signaling axis regulates metastatic potential of circulating tumor cells. However, modulation of immune cell infiltration into the tumor microenvironment and the elaboration of anti-tumor immunity, in particular by NK and CD8+ T cells may be involved. In fact, it was recently shown that loss of the S1P transporter Spns2, which is highly expressed in the endothelium, suppressed metastatic potential of circulating tumor cells in mouse models (36).

Using the genetic loss of function models of S1PR2 and S1PR3, either alone or in combination with S1PR1, we show that these two S1PRs compensate for the loss of S1PR1 in tumor vascular endothelium. This finding may be useful in the design of therapeutic approaches to enhance tumor vascular normalization.

The clinical relevance of our study is underscored by the finding that doxorubicin-induced tumor growth suppression is more effective in endothelial S1PR1 transgenic mice which show enhanced S1P signaling in the tumor vasculature.

In summary, our study shows that endothelial S1PR signaling is an important factor in tumor vascular phenotype that influences tumor progression, metastasis and chemotherapeutic efficacy. Strategies to enhance S1PR1 function in the tumor vasculature may potentiate the efficacy of cytotoxic and targeted anti-cancer therapies.

## Materials and Methods

### Mouse strains

Mice were housed in a temperature-controlled facility with a 12-h light/dark cycle, specific pathogen-free, in individual ventilated cages and provided food and water ad libitum. All animal experiments were approved by the Boston Children’s Hospital and Weill Cornell Medicine Institutional Animal Care and Use Committees. EC-specific *S1pr1* knockout mice (*S1pr1*^*f/f*^ Cdh5-Cre-ER^T2^; *S1pr1* ECKO) were generated as described (4, 20-22). EC-specific *S1pr2* knockout mice (*S1pr2*^*f/f*^ Cdh5-Cre-ER^T2^; *S1pr2* ECKO) were generated as follows. To create *S1pr2* conditional knockout targeting construct, *loxP* sites were inserted into the 5’ upstream of the first coding exon (exon 2), and downstream of the 3’ untranslated region. The following DNA segments were added immediately downstream of the 3’ *loxP* site: a *phosphoglycerate kinase* (*PGK*)-*Neo* cassette flanked by *frt* sites with a third *loxP* inserted immediately 5’ of the second *frt* site. *Xba I* site was inserted adjacent to second *loxP* site to facilitate analysis by Southern blotting. The targeting constructs were linearized and electroporated into C57Bl/6 embryonic stem (ES) cells, and after positive selection with G418, ES colonies were screened by Southern blot hybridization. Mice carrying the primary targeted alleles were crossed to germline *Flp* mice to excise the *frt-PGK-Neo-frt* cassette. They were then crossed to Cdh5-Cre-ER^T2^ mice to generate *S1pr2* ECKO mice. EC-specific *S1pr1-S1pr2* double knockout mice were generated by crossing *S1pr1* ECKO with *S1pr2* ECKO mice. EC-specific *S1pr1-S1pr2* double knockout mice in the *S1pr3*^-/-^ background were generated by crossing the *S1pr1* ECKO *- S1pr2* ECKO mice with *S1pr3*^-/-^ mice (6). *S1pr1*^*f/stop/f*^ was generated as described (4, 20) by knocking in the transgene into embryonic stem cells (ESCs) and crossed to Cdh5-Cre-ER^T2^ mice. Gene deletion or overexpression by the cre recombinase was achieved by intraperitoneal injection of tamoxifen (Sigma-Aldrich) (150 µg/gram of body weight/day) at 6 weeks of age for five consecutive days, and mice were allowed to recover for a week before being used for experiments. Littermates without the Cdh5–Cre–ERT2 gene were treated with tamoxifen in the same way and used as controls. S1P1-GFP reporter mice have been previously described (19). Briefly, mice expressing a transcriptional unit consisting of the S1PR1 carboxy terminus fused to tetracycline transcriptional activator (tTA) and ß-arrestin fused to tobacco etch virus (TEV) protease knocked in to the endogenous *S1pr1* locus were crossed with mice containing a histone–GFP reporter gene under the control of a tTA-responsive promoter. Mice expressing one allele of both transgenes were considered S1PR1 GFP signaling mice. Littermates expressing only the GFP allele without the S1P1 knock-in were considered controls (19). All genotyping was done by PCR using ear punch biopsies.

### Cell lines

Lewis lung carcinoma cells (LLC, ATCC-CRL-1642) used for subcutaneous injection and B16F10 cells (ATCC-CRL-6475) were grown in DMEM supplemented with 10% heat-inactivated fetal bovine serum (FBS). Both cell lines were tested with the IMPACTIII Rodent Pathogen Testing (IDEXX RADIL, University of Missouri) prior to experiments in mice.

### Tumor growth and drug administration

Lewis lung carcinomas (LLC) cells (5×10^5^ suspended in HBSS) were injected subcutaneously on both flanks into the indicated mice. 16 days later, tumors were harvested and analyzed further. For water content, tumors were weighed following harvest, dried overnight in a 60 degrees oven, and weighed again. Percent water content was calculated using the formula -((wet weight–dry weight)/wet weight)*100. Doxorubicin (Sigma-Aldrich) was administered at a final dose of 5 mg/kg body weight via intraperitoneal injection every other day, starting 8 days post tumor cell injection. Control animals were treated with the vehicle, HBSS. Tumor-bearing mice were sacrificed 22 days post LLC injection. B16F10 cells (10^6^ in HBSS) were injected intravenously in the tail vein into the indicated mice. 20 days later, mice were euthanized by CO2, perfused with 10 mL of PBS and lungs were harvested. Metastasis foci in lung tissue sections were counted under a microscope.

### Immunostaining and imaging

Tumors were fixed in 4% paraformaldehyde (PFA) in PBS at 4°C, washed in PBS and processed for paraffin or embedded in OCT compound (Tissue-Tek, Fisher) for frozen section. Paraffin sections (12µm) were stained either with Hematoxylin-Eosin, or with CD31 antibody (BD Pharmigen, clone MEC13.3) counterstained with hematoxylin, or Rabbit anti-*α*-SMA (Rabbit, Abcam) counterstained with methyl green (Vector Laboratories). Brightfield images were taken using Zeiss Axioskop2 microscope with an AxioCam digital camera (Zeiss). Cryosections (35 µm –tumors, 10 µm -lungs) were permeabilized with PBS - 0.1% Triton at RT for 30 minutes, then blocked with PBS containing 75 mM NaCl, 18 mM Na_3_ citrate, 2% FBS, 1% bovine serum albumin (BSA) and 0.05% Triton X-100. Tumor vasculature in frozen sections were obtained by the co-staining of rat anti-CD31 antibody, Cy3-conjugated anti-smooth muscle actin (*α*-SMA, Sigma) and or rabbit anti-Chondroitin Culfate Proteoglycan (NG2, Millipore). Endothelial S1PR1 expression was detected by a rabbit anti-S1PR1 (Santa Cruz, H60). Confocal images were taken using an Olympus FluoView FV10i or Zeiss LSM 800 with Airyscan confocal microscopes. The 3D reconstructions of z-stack (XY projection) images are shown. Image processing and quantification were performed by using Adobe Photoshop, ImageJ, or Fiji software (National Institutes of Health). CD31, SMA and NG2 positive immunofluorescent signals were subjected to threshold processing and areas occupied by their signal were quantified using the Fiji Software, as described previously (22). The vascular density was determined by normalizing CD31 positive area to the total area of the tumor. Branch point number was quantified on skeletonized CD31 signal and normalized to the total area of the tumor. Total vascular length was quantified on skeletonized CD31 signal. Mural cell coverage of the vessels by smooth muscle cell and pericytes was determined by normalizing the SMA or NG2 positive area to the CD31 positive area.

### Tumor vessel leakiness

Mice bearing LLC tumors were injected intravenously via tail vein 100 µl of a mix containing 0.25 mg of 70 kDa Tetramethylrhodamine-conjugated dextran (Molecular Probes) and 0.25 mg of 2000 kDa Fluorescein-conjugated dextran (Molecular Probes) in HBSS. After 90 minutes, mice were euthanized by CO_2_ and perfused with 10 mL of PBS. Tumors were harvested, fixed with 4% PFA at 4°C, washed in PBS and embedded on OCT. Frozen blocks were cut in cryosections of 50 µm and imaged with a Zeiss LSM 800 with Airyscan confocal microscope. Image processing and quantification was performed by using Adobe Photoshop, ImageJ, or Fiji software (National Institutes of Health). The percent of 70 kDa leakage was quantified using Image J software, by subtracting the 70 kDa-positive signal from the 2000 kDa-positive signal and measuring the remaining 70 kDa positive signal.

### Quantitation of tumor hypoxia

To determine the hypoxic area of the tumor vasculature, tumor bearing mice were injected via tail vein with the oxygen-sensitive compound Hypoxyprobe-1 (pimonidazole hydrochloride, Hypoxyprobe, Inc) at a dosage of 60 mg/kg body weight as previously described (4). After 90 min, mice were euthanized and perfused with 10 mL of PBS. The tumors were harvested, fixed in 4% PFA, sectioned, permeabilized, and stained with an anti-Hypoxyprobe-1 antibody and CD31 (BD Pharmigen). Image processing and quantification was performed using Adobe Photoshop, ImageJ, or Fiji software (National Institutes of Health). Hypoxyprobe-1 positive immunofluorescent signals were subjected to threshold processing and areas occupied by their signal were quantified by using the Fiji Software, and normalized to the total area of the tumor.

### Statistical Analysis

Statistical analyses were performed using GraphPad Prism software v.7.0. Two-tailed unpaired Mann-Whitney test or t test with Welch’s correction were used for direct comparison of two groups. Analysis of variance (ANOVA) followed by Sydak’s multiple comparisons test to compare all groups was used to determine significance between three or more test groups. All values reported are means ± SD. All animal experiments used randomization to treatment groups and blinded assessment.

## Acknowledgements

The authors thank Drs. Diane Bielenberg and Bruce Zetter for advice on metastatic experiments, Dr. David Zurakowski for biostatistics advice and Kristin Johnson for graphics assistance.

## Funding

This work was supported by NIH Grants HL89934, HL117798, and R35 HL135821 (to T.H.), a Fondation Leducq Transatlantic Network Grant (SphingoNet)(to T.H.); and a postdoctoral fellowship from the American Heart Association 18POST33990452 (to A.C.).

## Competing interests

The authors declare that they have no competing interests. T.H. discloses that he received research funding from ONO Pharmaceutical Corporation, consulted for Steptoe and Johnson, LLP, and Bridge Medicine Inc., and is an inventor on ApoM+HDL, S1P chaperones, and S1P receptor antagonists.

## References

1. Proia RL & Hla T (2015) Emerging biology of sphingosine-1-phosphate: its role in pathogenesis and therapy. J Clin Invest 125(4):1379–1387.

2. Liu Y, et al. (2000) Edg-1, the G protein-coupled receptor for sphingosine-1-phosphate, is essential for vascular maturation. J Clin Invest 106(8):951–961.

3. Paik JH, et al. (2004) Sphingosine 1-phosphate receptor regulation of N-cadherin mediates vascular stabilization. Genes Dev 18(19):2392–2403.

4. Jung B, et al. (2012) Flow-regulated endothelial S1P receptor-1 signaling sustains vascular development. Dev Cell 23(3):600–610.

5. Gaengel K, et al. (2012) The sphingosine-1-phosphate receptor S1PR1 restricts sprouting angiogenesis by regulating the interplay between VE-cadherin and VEGFR2. Dev Cell 23(3):587–599.

6. Kono M, et al. (2004) The sphingosine-1-phosphate receptors S1P1, S1P2, and S1P3 function coordinately during embryonic angiogenesis. J Biol Chem 279(28):29367–29373.

7. Mizugishi K, et al. (2005) Essential role for sphingosine kinases in neural and vascular development. Mol Cell Biol 25(24):11113–11121.

8. Xiong Y, Yang P, Proia RL, & Hla T (2014) Erythrocyte-derived sphingosine 1-phosphate is essential for vascular development. J Clin Invest 124(11):4823–4828.

9. Folkman J (2007) Angiogenesis: an organizing principle for drug discovery? Nat Rev Drug Discov 6(4):273–286.

10. Ferrara N (2010) Pathways mediating VEGF-independent tumor angiogenesis. Cytokine Growth Factor Rev 21(1):21–26.

11. Carmeliet P & Jain RK (2011) Principles and mechanisms of vessel normalization for cancer and other angiogenic diseases. Nat Rev Drug Discov 10(6):417–427.

12. Jain RK (2005) Normalization of tumor vasculature: an emerging concept in antiangiogenic therapy. Science 307(5706):58–62.

13. Huang Y, Goel S, Duda DG, Fukumura D, & Jain RK (2013) Vascular normalization as an emerging strategy to enhance cancer immunotherapy. Cancer Res 73(10):2943–2948.

14. Goel S, et al. (2011) Normalization of the vasculature for treatment of cancer and other diseases. Physiol Rev 91(3):1071–1121.

15. Chae SS, Paik JH, Furneaux H, & Hla T (2004) Requirement for sphingosine 1-phosphate receptor-1 in tumor angiogenesis demonstrated by in vivo RNA interference. J Clin Invest 114(8):1082–1089.

16. LaMontagne K, et al. (2006) Antagonism of sphingosine-1-phosphate receptors by FTY720 inhibits angiogenesis and tumor vascularization. Cancer Res 66(1):221–231.

17. Azuma H, et al. (2002) Marked prevention of tumor growth and metastasis by a novel immunosuppressive agent, FTY720, in mouse breast cancer models. Cancer Res 62(5):1410–1419.

18. Fischl AS, et al. (2019) Inhibition of Sphingosine Phosphate Receptor 1 Signaling Enhances the Efficacy of VEGF Receptor Inhibition. Mol Cancer Ther 18(4):856–867.

19. Kono M, et al. (2014) Sphingosine-1-phosphate receptor 1 reporter mice reveal receptor activation sites in vivo. J Clin Invest 124(5):2076–2086.

20. Blaho VA, et al. (2015) HDL-bound sphingosine-1-phosphate restrains lymphopoiesis and neuroinflammation. Nature 523(7560):342–346.

21. Galvani S, et al. (2015) HDL-bound sphingosine 1-phosphate acts as a biased agonist for the endothelial cell receptor S1P1 to limit vascular inflammation. Sci Signal 8(389):ra79.

22. Yanagida K, et al. (2017) Size-selective opening of the blood-brain barrier by targeting endothelial sphingosine 1-phosphate receptor 1. Proc Natl Acad Sci U S A 114(17):4531– 4536.

23. Helmlinger G, Yuan F, Dellian M, & Jain RK (1997) Interstitial pH and pO2 gradients in solid tumors in vivo: high-resolution measurements reveal a lack of correlation. Nat Med 3(2):177–182.

24. Chen J, et al. (2009) Suppression of retinal neovascularization by erythropoietin siRNA in a mouse model of proliferative retinopathy. Invest Ophthalmol Vis Sci 50(3):1329–1335.

25. Lee MJ, et al. (1999) Vascular endothelial cell adherens junction assembly and morphogenesis induced by sphingosine-1-phosphate. Cell 99(3):301–312.

26. Skoura A, et al. (2007) Essential role of sphingosine 1-phosphate receptor 2 in pathological angiogenesis of the mouse retina. J Clin Invest 117(9):2506–2516.

27. Sanchez T, et al. (2007) Induction of vascular permeability by the sphingosine-1-phosphate receptor-2 (S1P2R) and its downstream effectors ROCK and PTEN. Arterioscler Thromb Vasc Biol 27(6):1312–1318.

28. Adada M, Canals D, Hannun YA, & Obeid LM (2013) Sphingosine-1-phosphate receptor 2. FEBS J 280(24):6354–6366.

29. Windh RT, et al. (1999) Differential coupling of the sphingosine 1-phosphate receptors Edg-1, Edg-3, and H218/Edg-5 to the G(i), G(q), and G(12) families of heterotrimeric G proteins. J Biol Chem 274(39):27351–27358.

30. Hla T (2001) Sphingosine 1-phosphate receptors. Prostaglandins Other Lipid Mediat 64(1-4):135-142.

31. Sanchez T & Hla T (2004) Structural and functional characteristics of S1P receptors. J Cell Biochem 92(5):913–922.

32. Hisano Y & Hla T (2019) Bioactive lysolipids in cancer and angiogenesis. Pharmacol Ther 193:91–98.

33. Pitulescu ME, Schmidt I, Benedito R, & Adams RH (2010) Inducible gene targeting in the neonatal vasculature and analysis of retinal angiogenesis in mice. Nat Protoc 5(9):1518– 1534.

34. Cantelmo AR, et al. (2016) Inhibition of the Glycolytic Activator PFKFB3 in Endothelium Induces Tumor Vessel Normalization, Impairs Metastasis, and Improves Chemotherapy. Cancer Cell 30(6):968–985.

35. Gaengel K, Genove G, Armulik A, & Betsholtz C (2009) Endothelial-mural cell signaling in vascular development and angiogenesis. Arterioscler Thromb Vasc Biol 29(5):630–638.

36. van der Weyden L, et al. (2017) Genome-wide in vivo screen identifies novel host regulators of metastatic colonization. Nature 541(7636):233–236.

